# Species-Specific Responses Drive Browsing Impacts on Physiological and Functional Traits in *Quercus agrifolia* and *Umbellularia californica*

**DOI:** 10.1101/2023.06.01.543299

**Authors:** Hugh E. Leonard, Mary Ciambrone, Jarmila Pittermann

**Affiliations:** Department of Ecology and Evolutionary Biology, University of California, Santa Cruz, Santa Cruz, California, United States of America

## Abstract

Herbivory is a fundamental ecological force in the evolution of plant physiological, morphological, and chemical attributes. In this study, we explored how browsing pressure by local deer populations affected leaf form and function in two California native tree species, coast live oak (*Quercus agrifolia*) and bay laurel (*Umbellularia californica*). Specifically, we investigated how leaf and stem vascular attributes shifted between browsed and non-browsed zones of each species and tested for differences in trait coordination as well as stem-leaf function. Browsing significantly altered traits such as leaf to phloem ratios and leaf area, but we observed few meaningful differences in leaf and stem anatomy between browsed and non-browsed material. We discuss these results in the context of such additional ecological factors and explore future research considerations for investigating leaf and stem vascular trait development with herbivore presence.

## Introduction

Herbivory is a well-documented biological interaction influencing the ecology of natural systems [1-4]. Herbivores are ecosystem engineers because they can alter ecosystem structure and function through consumer activities and physical presence [5-7]. As primary consumers, herbivores impact ecosystem function by reducing aboveground biomass, altering resource allocation during competition with other plants, and reducing recruitment of new shoots and seedlings [8, 9]. A notable example of this is the re-introduction of wolves into Yellowstone National Park, which decreased elk presence in willow thickets and led to an increased aspen biomass [8]. This and other examples demonstrate how herbivore presence and density can transform the plant community’s physical structure and species composition, all of which can alter ecosystem function.

Herbivory can dramatically alter a plant’s morphometric traits at the level of individual plants. The most direct impacts arise from the alteration of resource allocation throughout the plant [10, 11, 12]. A notable example is the atypical growth that occurs when browsers remove the softer leaves and buds of the apical meristem region. Terminal shoots produce auxin (indole-3-acetic acid), with bud removal lowering hormone action and reducing apical dominance [13-15]. Through browsing, herbivores can stimulate plants to grow branches with shortened internode distances and compact foliage that produce smaller leaf surface areas and thinner leaves [16, 17]. Thinner and smaller leaves may result from reduced carbon resources available for growth, but this response may also be adaptive. For example, reduction in leaf area and mass can be viewed an adaptation to herbivory by reducing the carbon cost to the plant in production of vulnerable leaves, in comparison to the upregulation of the more carbon-costly secondary compounds that reduce leaf palatability [3]. However, changes in leaf vascular structure and development may be the downstream consequence of reduced leaf area and carbon investment, and this may impact overall shoot water transport and photosynthetic capacity [18, but see also 19-22].

Previous work on herbivory’s impact on plant performance has shown that long-term browsing reduces vessel density, stem hydraulic conductivity, and Huber values [18]. However, how herbivory affects the coordination between plant leaf and stem vascular tissue growth remains unknown. Specifically, it is unclear how browsing affects leaf cell and tissue structure, especially stomatal and epidermal cell traits critical to leaf water relations [20], and whether these changes could lead to permanent alterations in the structure and function of leaves and stems. Many tree and shrub species experience intensive browsing at early life stages until growing high enough to escape this ‘browse trap’ [6], resulting in effectively two growth habits in one individual: browsed and non-browsed (Fig 1A, B). A targeted investigation of leaf and stem anatomical and functional trait responses to browsing, conducted across multiple taxa, is necessary to understand herbivory’s direct impacts on morphology and hydraulic function.

**Figure 1.**
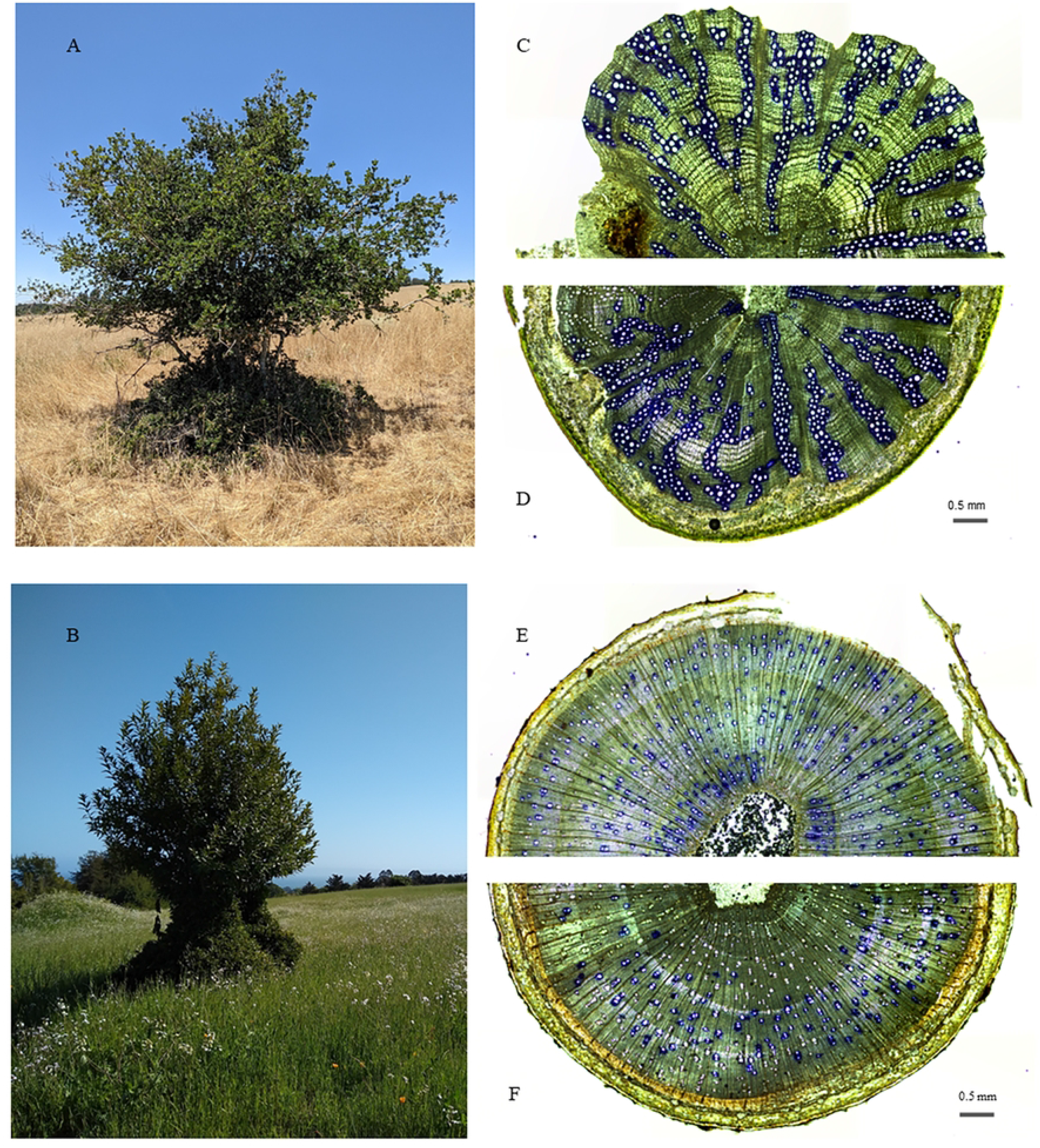
Morphology and stem anatomy of Q. agrifolia (A, C, D) and U. californica (B, E, F). Image A and B show tree morphology and topiary structure from deer browsing. Images C-F show functional xylem (stained blue) and general stem anatomy for non-browsed (C, E) and browsed (D, F) zones of each species.

To this end, we analyzed the effect of browsing activity of black-tailed deer (*Odocoileus hemionus columbianus*) on plant performance using leaf and vascular traits for two woody evergreen species, coast live oak (*Quercus agrifolia)*, a ring-porous species, and bay laurel (*Umbellularia californica)*, which has diffuse-porous xylem. We sought to understand how responses to herbivory can a) alter leaf and stem morphology and anatomy and b) what effect(s) these changes can have on leaf and stem vascular function. Both tree species occur in the same habitat and are exposed to the same deer browsing pressure, allowing cross-species comparisons in a natural setting. Significantly for this study, both tree species incurred deer browsing only between ground level and approximately 1.5 meters in height, above which the trees had no browsing impact. Leaf traits including leaf mass per area (LMA), leaf vein density, stomatal and leaf epidermal cell density and size, and stomatal conductance were collected, as well as stem vascular traits including midday water potential, stem functional xylem, xylem conduit size, and density, and phloem tissue width in browsed and non-browsed zones for each species. We hypothesized that browsing would result in smaller leaves with smaller and fewer stomata, in coordination with reduced xylem and phloem vessel size and densities. Ultimately, we sought to clarify the mechanism of herbivory as a driver of vascular development and function.

## Methods

### Site Description

The study was executed at the University of California, Santa Cruz. The university is located on the southwestern edge of the Santa Cruz Mountains and experiences a Mediterranean climate with warm, dry summers and cool, moist winters. The average annual rainfall is 79 cm, and many summer days are characterized by afternoon and evening fog events [23]. Coast live oak specimens for this study occupied an open grassland area with full sun exposure. Bay laurel specimens were located within the same open grassland and were adjacent to a mixed evergreen forest edge. The surrounding habitat is primarily open and composed of low-growing, herbaceous species of grass and forbs (Fig 1A, B).

### Experimental Design

Coast live oak and bay laurel species were selected for this study from field observations of heavy browsing from the local black-tailed deer population that resulted in a strongly topiary growth habit (Fig 1A, B). The altered shape of these trees presented a natural experiment for investigating the physiological impact of browsing on leaf and stem tissues within the same tree. All samples and measurements were conducted on sun-exposed branches selected from a browsed height of less than 1.5 meters (hereafter called the browsed zone) and non-browsed height of 1.5-2 meters (hereafter called the non-browsed zone) of each tree. Leaf and stem material were collected in the field during late spring on recently developed, fully mature growth with leaves undamaged by browsing and transported to the lab for processing. Midday water potentials and stomatal conductance measurements were conducted in the field during the beginning and end of the dry season (June-October) from browsed and non-browsed zones to capture functional impacts on seasonal hydraulic capacity from browsing. All field data was collected between January 2020 and December 2021.

### Leaf Traits

#### Collection of LMA

12 undamaged leaves were collected from both browsed and non-browsed zones of ten trees per species, for a total of 240 leaves per species (n=120 for browsed and non-browsed zones). Collected leaves were wrapped in moistened paper towels and placed in plastic bags for transport to the lab. Leaves were digitally scanned (HP Photosmart C6380 All-In-One Printer/Scanner; Hewlett-Packard, Palo Alto, CA, USA) and images were processed using ImageJ software for leaf area (cm2; http://rsbweb.nih.gov/ij/index.html). After scanning, leaves were oven-dried for three days at 70 degrees Celsius and dry mass was recorded [18]. Each leaf was weighed on a digital balance (grams; Acculab ALC series; Sartorius Group, Germany) avoiding air exposure until mass was recorded. LMA was calculated as the ratio of dry leaf mass to leaf area [24].

#### Vein Density

Three undamaged leaves were collected from browsed and non-browsed zones of ten coast live oak and ten bay laurel trees, resulting in a total of 60 leaves per species (n=30 for browsed and non-browsed zones). Biopsy punches were used to generate three 4 mm diameter samples per leaf (RoyalTek biopsy punch; Surgical Design Inc, Lorton, VA, USA), selecting regions devoid of primary veins. Each leaf sample was treated with a solution of 1.0 M KOH in deionized water for pigment clearing. After 8 to 14 days, depending on the specific leaf samples and clearing rates, samples were removed and washed with deionized water before staining with toluidine blue. Stained leaf sections were mounted in glycerol and photographed under a 4X objective on a compound light microscope (Leica model DM6 B; Leica Microsystems, Germany) and analyzed for vein density using ImageJ software. Vein density for each species was calculated as the total vein length per leaf area averaged across each biopsy punch [25].

#### Stomatal Traits

Stomatal measurements were made using the same mounted leaf samples used for vein density analysis. Stomatal density was captured under a 10X objective and measured as the number of stomata per square millimeter of leaf area. For stomatal anatomy measurements, stomata were selected from the same region used for density measurements and photographed under 20X and 40X objectives. Each stoma within a region was numbered, and the length and width of 15 randomly selected stomata per leaf were measured (n=150 for browsed and non-browsed zones, per species). Stomatal size was approximated using stomatal length multiplied by stomatal width [26].

#### Epidermal Cell Traits

Epidermal measurements, specifically pavement cells [27], were made on the same mounted leaf samples used for vein density measurements. Fifteen pavement cells per leaf were imaged under 20X and 40X microscope objectives (n=150 for browsed and non-browsed zones, per species). Selection of pavement cells was established by outlining a region of leaf and selecting cells from four corners of the viewing field and one cell from the center. Pavement cell area was measured manually by tracing the outline of each cell using ImageJ software and recording the area enclosed.

#### Stomatal Conductance and Midday Water Potential

To assess differences in water stress between browsed and non-browsed regions, mid-day water potential (Ψ_mid_) and stomatal conductance were collected from six coast live oak and bay laurel trees (n=6 trees per species). Measurements were collected during June of 2021 prior to the onset of the summer dry period and again in October 2021 at the end of the summer dry season. Measurements were obtained from exposed leaves of at least one year of age and collected between 10 am and 3 pm to capture maximum leaf transpiration and xylem water tension. Stomatal conductance was collected using a portable porometer (SC-1 leaf porometer; Decagon Devices, Pullman, WA, USA) on three separate leaves, and Ψ_mid_ was collected on the same branchlets using a Scholander pressure chamber [28].

### Stem Traits

#### Stem Selection and Age

Stem traits were obtained from six trees per species (n=6 per species), with one branch selected from the browsed zone and one from the non-browsed zone of each tree. Branch length and internode distance varied between browsed and non-browsed zones due to deer browsing, therefore direct observation could not be reliably used to determine branch age. To ensure that age was not a confounding factor in stem structure and function, each branch was confirmed to be between two and four years of age based on the number of growth rings present after cutting. Branches were covered in dark plastic bags before dawn for a minimum of 15 minutes to allow water potential within the xylem water column to equilibrate to the atmospheric conditions and reduce tension on the xylem water column [18, 29]. Each branch was then cut and quickly submerged under water before cutting an additional 3cm of stem to minimize the introduction of air into conductive xylem vessels. Sample branches were then transported within three hours to the lab for processing, with the cut ends kept submerged to avoid introduction of air into xylem vessels and embolism formation.

#### Functional Xylem

To identify functional xylem vessels, protocols were followed as outlined by Jacobsen et al [30], imposing a negative pressure of 2kPa on 12 cm distal stem portions to draw crystal violet stain into the functional vessels of the stem. This pressure was applied for five minutes, after which segments were removed from the vacuum system and deionized water was used to flush excess stain from the segments to prevent false positives in conductive vessel identification. All segments were then frozen immediately to avoid diffusion of stain into non-conductive tissues, and frozen segments were used to generate cross-sections from the center of each segment using a sledge microtome. Freezing the samples avoided excessive sample fracturing during sectioning.

Prepared cross-sections were mounted on slides with glycerol and photographed under 4X objective. To assess conductive vessel density, cross-sections were partitioned using ImageJ into three 33 degree wedges (± 1 degree) extending from pith the bark and arranged equidistant around the cross section when possible. Within each wedge, all conductive vessels were counted and diameters were taken for both the long and short widths. These diameters were then used to compute the hydraulic diameter for each vessel and extrapolated as the average hydraulic diameter for each cross section [31].

#### Phloem Width

Phloem data was collected on the same cross sections used for xylem analysis. Phloem width was determined from the average of four width measurements collected from each cross section and separated by 90 degrees when possible. Phloem area was calculated as the difference in stem area minus bark and xylem tissue, and measurements were made using the assumption of a circular area (πr^2^).

### Data Analysis

All statistical analysis was conducted using R 3.6.3 [32]. To compare differences in browsing effects for each species, data was averaged across specimen replicates to generate averages for each tree. Two-tailed t-tests were used to compare differences between browsed and non-browsed zones per tree. Linear regression models were made using the lm() function to assess trait coordination and response to herbivory within browsed and non-browsed zones. Results were considered significant if p < 0.05.

## Results

### Leaf Morphometric and Anatomical Traits

Leaves were generally smaller and had lower LMA in browsed zones, although the differences were inconsistent across species (Table 1, Fig 2). For example, leaves were approximately 57% smaller (p=0.00607) in browsed bay laurel compared to non-browsed zones. However, LMA remained largely invariable in browsed and non-browsed zones. In oak, by contrast, leaf mass per area was 11% lower (Table 1, p=0.035) in browsed zones while leaf area was not statistically different from non-browsed zones, suggesting that continual browsing reduces the available resources needed to construct robust new leaves. (Fig 2B). Taken together, this suggests that both leaf size and LMA vary as a function of species-specific responses.

**Figure 2.**
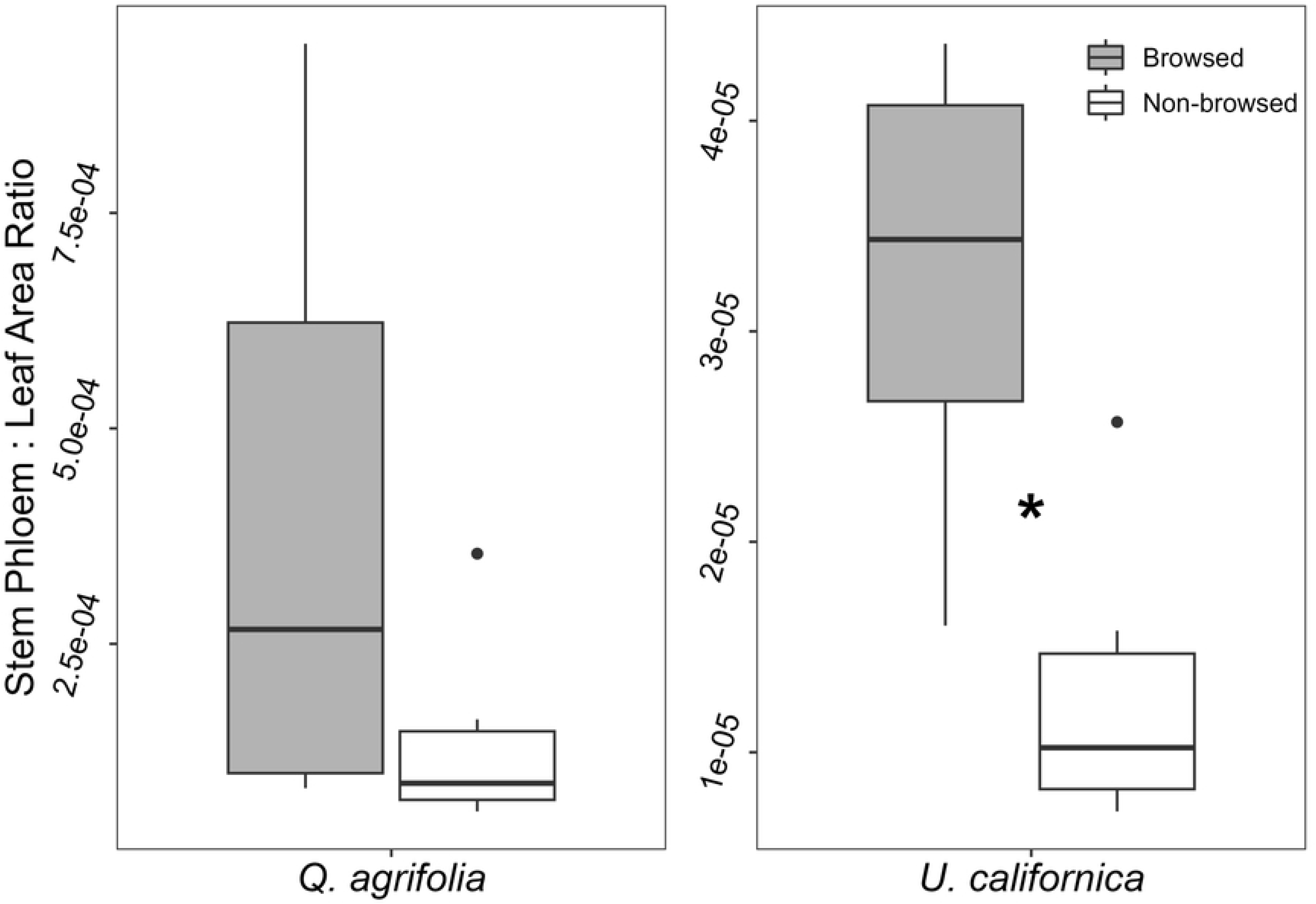
Comparison of browsed and non-browsed leaf traits. Boxplots of browsed and non-browsed (A) leaf area, (B) conductive xylem area, (C) leaf mass per area, and (D) conductive vessel density. *U. californica* showed a significantly larger leaf area for non-browsed zones (p = 0.00607), and *Q. agrifolia* had a significantly greater LMA for non-browsed zones compared to browsed zones (p = 0.049). *U. californica* browsed zones contained significantly less conductive xylem (p = 0.0099) and had a lower density of conductive vessels (p = 0.0314) compared to non-browsed zones.

**Table 1.** Comparison of Physiological Traits Between Browsed and Non-browsed Zones.

| *Q. agrifolia* |  |  |  |
| --- | --- | --- | --- |
| Trait | Browsed | Non-browsed | p-value |
| Internode Length (cm) | 2.24 ± 0.98 | 11.33 ± 3.30 | **5.2e-07** |
| Branch Length (cm) | 30.4 ± 16.64 | 55.15 ± 11.11 | **0.0148** |
| LMA (g m<sup>-2</sup>) | 128.48 ± 19.33 | 144.31 ± 13.36 | **0.049** |
| Leaf Area (cm<sup>2</sup>) | 299.021 ± 219.70 | 628.46 ± 406.13 | 0.12 |
| Stomatal Size (um<sup>2</sup>) | 593.94 ± 75.98 | 635.38 ± 43.66 | 0.157 |
| Huber Value (functional xylem/leaf area) | 3.33e-04 ± 1.92e-04 | 1.77e-04 ± 1.71e-04 | 0.17 |
| Pavement Cell Size (um<sup>2</sup>) | 215.72 ± 42.29 | 191.14 ± 38.02 | 0.189 |
| Mean Phloem Width (cm) | 0.035 ± 7.09e-03 | 0.040 ± 2.72e-03 | 0.218 |
| Conductive Vessel Density (m<sup>-2</sup>) | 3.59e+07 ± 9.30e+06 | 2.95e+07 ± 7.61e+06 | 0.225 |
| Vein Density (per cm<sup>2</sup>) | 819 ± 186 | 947 ± 266 | 0.23 |
| Conductive Mean Conduit Diameter (um) | 33.84 ± 4.61 | 37.01 ± 4.76 | 0.268 |
| Conductive Hydraulic Diameter (um) | 32.50 ± 4.51 | 35.58 ± 4.76 | 0.277 |
| Stomatal Density (cm<sup>-2</sup>) | 41950 ± 8247 | 46180 ± 10169 | 0.321 |
| Phloem to Xylem Width Ratio | 0.17 ± 0.040 | 0.18 ± 0.026 | 0.467 |
| Stem Areas (cm<sup>2</sup>) | 0.54 ± 0.14 | 0.56 ± 0.14 | 0.779 |
| Stomatal Conductance (mmol m<sup>-2</sup> s<sup>-1</sup>) | 474.96 ± 58.38 | 466.47 ± 64.72 | 0.787 |
| Midday Water Potential (MPa) | 1.23 ± 0.56 | 1.28 ± 0.36 | 0.826 |
| Conductive Xylem to Stem Area Ratio | 0.12 ± 0.033 | 0.12 ± 0.048 | 0.955 |
| Phloem Area (cm<sup>2</sup>) | 0.056 ± 0.010 | 0.056 ± 0.013 | 0.973 |
| Mean Xylem Width (cm) | $0.22 \pm 0.052$ | $0.22 \pm 0.019$ | 0.992 |
| Conductive Xylem area (cm<sup>2</sup>) | $0.067 \pm 0.028$ | $0.066 \pm 0.015$ | 0.9995 |
| *U. californica* |  |  |  |
| **Trait** | **Browsed** | **Non-browsed** | **p-value** |
| Internode Length (cm) | $3.15 \pm 1.62$ | $23.41 \pm 7.59$ | **1.03e-6** |
| Leaf Area (cm<sup>2</sup>) | $1139.44 \pm 796.59$ | $2633.83 \pm 684.54$ | **0.00607** |
| Conductive Xylem area (cm<sup>2</sup>) | $0.15 \pm 0.083$ | $0.31 \pm 0.090$ | **0.0099** |
| Branch Length (cm) | $47.10 \pm 21.29$ | $91.53 \pm 27.48$ | **0.0115** |
| Conductive Vessel Density (m<sup>-2</sup>) | $2.02e+07 \pm 9.85e+06$ | $3.43e+07 \pm 6.24e+06$ | **0.0314** |
| Stem Areas (cm<sup>2</sup>) | $0.41 \pm 0.095$ | $0.62 \pm 0.20$ | 0.0582 |
| Conductive Xylem to Stem Area Ratio | $0.35 \pm 0.15$ | $0.50 \pm 0.045$ | 0.0582 |
| Mean Xylem Width (cm) | $0.21 \pm 0.040$ | $0.25 \pm 0.033$ | 0.0741 |
| Phloem to Xylem Width Ratio | $0.10 \pm 0.032$ | $0.076 \pm 0.014$ | 0.146 |
| Huber Value (functional xylem/leaf area) | $1.48e-04 \pm 4.20e-05$ | $1.18e-04 \pm 2.30e-05$ | 0.158 |
| Conductive Hydraulic Diameter (um) | $27.26 \pm 5.83$ | $31.07 \pm 0.80$ | 0.218 |
| Conductive Mean Conduit Diameter (um) | $28.86 \pm 6.15$ | $32.76 \pm 0.76$ | 0.23 |
| Midday Water Potential (MPa) | $1.12 \pm 0.37$ | $0.98 \pm 0.37$ | 0.419 |
| Pavement Cell Size (um<sup>2</sup>) | $534.72 \pm 153.45$ | $490.79 \pm 116.12$ | 0.48 |
| LMA (g m<sup>-2</sup>) | $111.08 \pm 15.99$ | $115.92 \pm 21.57$ | 0.576 |
| Phloem Area (cm<sup>2</sup>) | $0.028 \pm 0.011$ | $0.031 \pm 0.012$ | 0.604 |
| Stomatal Size (um<sup>2</sup>) | $419.31 \pm 71.57$ | $405.11 \pm 63.68$ | 0.645 |
| Stomatal Density (cm<sup>-2</sup>) | $21391 \pm 6648$ | $20397 \pm 4879$ | 0.708 |
| Mean Phloem Width (cm) | $0.02 \pm 6.22\text{e-}03$ | $0.019 \pm 4.80\text{e-}03$ | 0.712 |
| Stomatal Conductance ($\text{mmol m}^{-2} \text{s}^{-1}$) | $179.09 \pm 61.96$ | $168.80 \pm 63.42$ | 0.718 |
| Vein Density (per $\text{cm}^2$) | $458 \pm 140$ | $434 \pm 159$ | 0.725 |

Trait means, ± standard deviation, with Welch's t-test outputs comparing browsed and non-browsed leaf and stem tissue for *Q. agrifolia* and *U. californica*.

Vein densities for coast live oak and bay laurel were similar for leaves from browsed and non-browsed zones (Table 1). Likewise, both species’ size and density of leaf stomatal and pavement cells did not differ between browsed and non-browsed zones. However, we did notice interesting relationships between leaf traits in both species. For example, an increase in pavement cell size was associated with lower stomatal density in both species (Table 2). However, this was observed in leaves from non-browsed zones in oak (p = 0.02) and browsed zones in bay laurel (p = 0.003). Furthermore, we examined the relationship between leaf area and leaf venation in both species and found that only coast live oak leaves from browsed zones had higher densities of veins with increasing stomatal density (p = 0.041). This suggests that leaf development in response to herbivory is inconsistent and can vary in different species.

**Table 2:** Significant functionally coordinated traits across species.

| Species | Type | Independent Trait | Dependent Trait | p-value | R<sup>2</sup> | Coefficient Estimate (slope) |
| --- | --- | --- | --- | --- | --- | --- |
| *Q. agrifolia* | Non-browsed | Total Leaf Area | Conductive Hydraulic Diameter | **0.00480** | 0.86 | 0.81 |
| *Q. agrifolia* | Non-browsed | Mean Leaf Size | Conductive Hydraulic Diameter | **0.0205** | 0.72 | 2.33 |
| *Q. agrifolia* | Non-browsed | Pavement Cell Size | Stomatal Density | **0.0201** | 0.45 | -1.92 |
| *Q. agrifolia* | Browsed | Mean Leaf Size | Conductive Hydraulic Diameter | **0.0241** | 0.70 | 3.74 |
| *U. californica* | Non-browsed | Mean Leaf Size | Conductive Xylem Area | **0.0477** | 0.58 | 0.014 |
| *U. californica* | Browsed | Total Leaf Area | Conductive Xylem Area | **0.00386** | 0.86 | 0.0098 |
| *U. californica* | Browsed | Pavement Cell Size | Stomatal Density | **0.0033** | 0.64 | -0.36 |

Regression analysis results from linear model, showing significantly correlated traits ( $p < 0.05$ ). Trait interactions varied substantially across *Q. agrifolia* and *U. californica*, suggesting a strong species-specific response to herbivory.

Taken together, leaf trait responses to herbivory were variable across each species. While leaves from browsed zones were, on average, half the size of the non-browsed zone leaves, leaf tissue investment varied by species, with lower LMA in the browsed zones of oak and no difference between bay laurel zones. Anatomical traits related to pavement cell size and density, stomatal size and density, and vein density were also unchanged in browsed zones relative to non-browsed zones. However, the relationships between these traits varied at a species-specific level to herbivory.

### Water Potential and Stomatal Conductance

Neither species exhibited differences in Ψ_mid_ or stomatal conductance in browsed and non-browsed zones during the sampling period, although both midday water potential and stomatal conductance were lower toward the end of the growing season (Table 3). This suggests that the water relations of the trees achieved some kind of equilibrium between the browsed and non-browsed regions. Bay laurel’s water potential, however, was significantly lower than coast live oak at the end of the season; this species may not be as deep-rooted as the coast live oaks.

**Table 3:** Seasonal midday water potential and stomatal conductance for coast live oak and bay laurel.

| Quercus |  |  |  |  |  |
| --- | --- | --- | --- | --- | --- |
| Species | Zone | Trait | Early summer | Late summer | p-value |
| Q. agrifolia | Browsed | Stomatal Conductance (mmol m<sup>-2</sup> s<sup>-1</sup>) | 470.94 ± 61.86 | 376.21 ± 52.52 | 0.00965 |
|  |  | Midday Water Potential (MPa) | -1.18 ± -0.58 | -1.0037 ± -0.34 | 0.75 |
|  | Non-browsed | Stomatal Conductance (mmol m<sup>-2</sup> s<sup>-1</sup>) | 467.57 ± 69.83 | 414.71 ± 75.11 | 0.198 |
|  |  | Midday Water Potential (MPa) | -1.23 ± -0.35 | -1.20 ± 0.32 | 0.859 |
| Umbellularia |  |  |  |  |  |
| Species | Zone | Trait | Early summer | Late summer | p-value |
| U. californica | Browsed | Stomatal Conductance (mmol m<sup>-2</sup> s<sup>-1</sup>) | 153.59 ± 44.00 | 99.96 ± 48.73 | 0.0518 |
|  |  | Midday Water Potential (MPa) | -1.16 ± -0.42 | -2.30 ± -0.82 | 0.00913 |
|  | Non-browsed | Stomatal Conductance (mmol m<sup>-2</sup> s<sup>-1</sup>) | 157.74 ± 72.13 | 101.55 ± 51.92 | 0.156 |
|  |  | Midday Water Potential (MPa) | -0.94 ± -0.41 | -2.49 ± -0.73 | 0.00198 |

Trait means, ± standard deviation, with p-values representing changes in trait values from early to late summer.

### Stem Morphometric and Anatomical Traits

Browsed zones extended 1.5 meters from the tree’s base, with leaves above that height presumably being out of reach for the deer. The perpetual nibbling created an obvious topiary architecture due to reduced growth at the base of the trees. Consequently, browsed branch length was substantially shorter in coast live oak stems than in non-browsed stems (Table 1; p = 0.0148), and a similar outcome was observed in U. californica (p = 0.0115). In addition to shorter branches, branch internode length was significantly shorter for browsed branches in coast live oak (p = 5.2e-07) and in bay laurel (Table 1; p = 1.03e-06), producing a condensed canopy of foliage under browsing pressure.

### Xylem Anatomy and Functional Traits

In coast live oak, no traits related to xylem or phloem function appeared to differ between browsed and non-browsed stems significantly. By contrast, bay laurel showed a significant 48% reduction in conductive xylem area in stems from the browsed zone (Fig 2B, Table 1; p = 0.0099). Due to the concurrent reduction in both leaf and conductive xylem areas for browsed stems, bay laurel Huber Values were similar for both browsed and non-browsed stems (1.48e-04 ± 4.20e-05 browsed and 1.18e-04 ± 2.30e-05 non-browsed, p = 0.158). However, the decrease in leaf area did not lead to a decrease in stem phloem area in browsed stems (Table 1, Fig 3; p = 0.604), suggesting a disconnect between phloem production and leaf area trait function.

**Figure 3.**
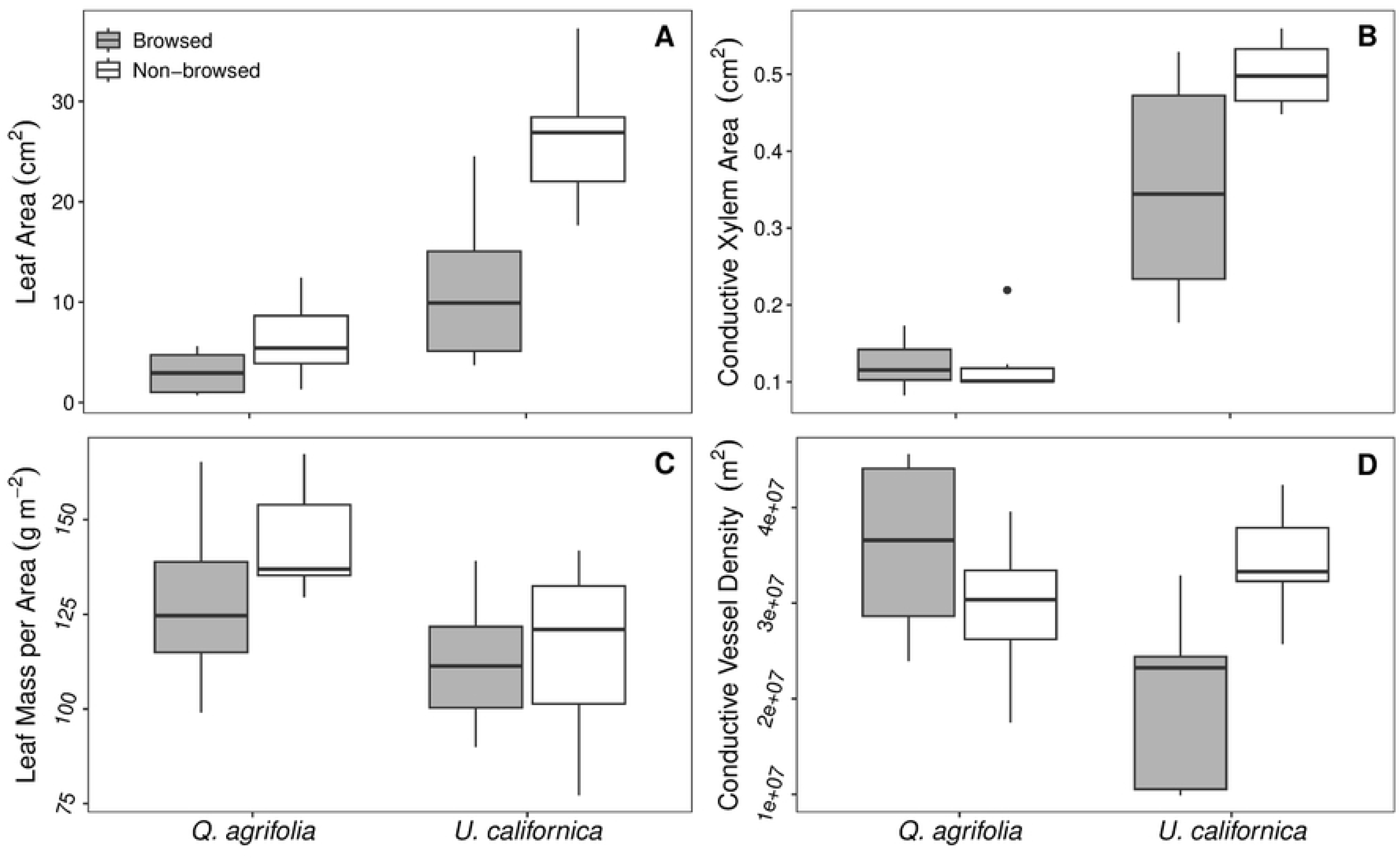
Ratio of stem phloem area to leaf area for coast live oak and bay laurel. Browsed zones of bay laurel had a significantly lower leaf area compared to non-browsed zones (p = 0.0046) but no reduction in phloem area. Significant differences are represented by asterisks.

## Discussion

Within this study, we sought to answer questions concerning browsing impacts on the development and function of leaf and stem vascular traits, particularly how browsing may alter tissue form and function within an individual tree. The results suggest that browsing uniquely impacts each species’ leaf and stem vascular function, but several traits vary consistently in direct response to browsing.

### Leaf Traits

LMA declined significantly in browsed zones of coast live oak but not in bay laurel, while mean leaf area declined in browsed zones of bay laurel but not in browsed coast live oak. Interestingly, each species has a specific trait response to browsing; however, reductions in leaf size and mass correspond with previous studies of herbivore impacts on leaf development [33, 16]. Changes in leaf size and LMA could result from browsing effects and the alteration of auxin production in terminal buds. Previous research has shown that repeated wounding increases auxin production within the surrounding plant tissues [34]. Higher auxin levels signal the release of jasmonic acid hormones (JAs) for defense. Increased JA levels have been correlated with stunted growth patterns in plants, primarily in the form of reduced tissue cell numbers [34, 35]. Potential ramifications from the reduction in leaf area and thickness include changes in water transport throughout the leaf and leaf water loss. Thinner leaves allow a decreased resistance to water exiting leaf veins and interacting with stomatal pores [36, 19], which, all other factors being equal, could lead to increased rates of water loss in browsed leaves.

In other aspects of leaf structure, the effects of herbivory become more nuanced. We expected herbivory to produce "cheaper" leaves with reduced nutrient allocation to browsed zones, resulting in lower vein density and a corresponding reduction in stomatal density. Instead, we found that both species’ vein and stomatal density were invariable across browsed and non-browsed leaves. This suggests that leaf vascular structure is not as strongly impacted by herbivory as we predicted. Selection pressure may not be so strong as to select for novel traits in browsed leaves, especially in trees. Additionally, there was no difference in stomatal conductance rates between zones, which correlates with the lack of stomatal size or density changes. Past research has shown that simulated defoliation can lead to nearly doubling stomatal conductance rates for defoliated stems compared to control treatments, likely due to an increased root-to-shoot ratio [29, but see 37]. In our study, using mature trees that maintained intact foliage after browsing may have obscured the shifts in stomatal conductance that were observed in previous studies on smaller individuals.

Variation in epidermal pavement and stomatal cell size was of particular interest in our study, as research has shown that epidermal cell size impacts leaf water loss [20]. Our initial hypothesis was that herbivory would reduce stomatal size and density due to reduced demand for water in response to lower leaf area. We further speculated that smaller pavement cells would arise as a consequence of developmental coordination. However, browsing had no effect on pavement or stomatal cell sizes or densities for either species. By contrast, we did observe coordination between increasing pavement cell size and decreasing stomatal density, but only in leaves collected from non-browsed zones of coast live oak and browsed zones of bay laurel. It is unclear why browsing would lead to opposite effects in coordination between these two traits, but possible explanations include species-specific reallocation of resources in browsed leaves or alteration of hormone production and regulation in developing leaf tissue when browsing occurs [38, 39].

### Vascular traits

Browsing effects varied between species and were species dependent. In coast live oak, the browsed and non-browsed tissues showed no significant differences in the vascular traits investigated. While an apparent reduction in internode length was observed in the field and for sampled branches, this difference did not appear to translate toward altering vascular traits within the stem. In bay laurel, however, several traits did show a significant difference between treatment types.

For bay laurel, conductive stem vessel densities and conductive xylem areas were significantly lower in browsed zones than in non-browsed. This is an intuitive result of browsing pressure, as breakage of vessels during browsing leads to air entry and embolism formation [40]. It is interesting to note the difference with coast live oak, as similar browsing damage to stems appeared across both species. This shift between species could be attributed to the differences in the vascular arrangement within stems, with coast live oak exhibiting a clustered "bundle" of vessels compared to the diffuse-porous structure in bay laurel (Fig 1C-E). This vascular arrangement could enhance redundancy to embolism formation within coast live oak. Additionally, coast live oak may already be so well adapted to drought stress that herbivore browsing may not significantly increase vascular stress from cavitation and embolism formation. Being extremely drought tolerant, coast live oak may inherently possess the hydraulic adaptations required for herbivore browsing resistance than bay laurel [41].

While bay laurel reduced conductive vessel density within browsed zones, it did not show a corresponding reduction in stomatal conductance or Ψ_mid_ for browsed stems, nor did it show altered stem vessel dimensions. Rather, the reduction in leaf area appears to be the primary driver of the reduction in conductive stem vessels. This response can be explained by reduced water supply to developing leaves, reducing leaf expansion [42-43]. Smaller leaves require less water to maintain turgor, allowing similar stomatal conductance rates as larger leaves and similar water potentials as stems from non-browsed zones. An important consequence of this reduced leaf size could be a net reduction in photosynthate produced, leading to a potential reduction in biomass accumulation for browsed zones [44].

### Trait Coordination

While the response of leaf and stem traits was inconsistent in browsed and non-browsed coast live oak, the interaction between traits was much more variable. Linear regression analysis between paired traits in browsed and non-browsed zones of each species showed significant correlations between traits, but the relationship was often varied both within and across species (Table 2). For example, non-browsed coast live oak zones showed positive correlations of conductive hydraulic diameter with both total leaf area and average leaf size, while browsed zones showed a positive correlation with only conductive hydraulic diameter and total leaf area. For bay laurel, conductive xylem area was positively correlated with mean leaf size in non-browsed zones and total leaf area in browsed zones. Interestingly, each species has a unique response to decreasing leaf size; coast live oak reduces the conductive hydraulic diameter, while bay laurel reduces the conductive xylem area. While effectively accomplishing the same net result, a reduced investment in water conducting tissue, the anatomical traits producing this change appear unique to each species. Although we did not investigate the species-specific mechanisms underlying these developmental differences, a likely factor would be the reduction of auxin regulation in browsed zones, which actively regulates cell differentiation and arrangement in developing stem and leaf tissue [45, 46, 47].

## Conclusion

The findings presented in this paper suggest that herbivory can be a complex moderator of leaf and stem adaptations. The ability of plants to respond to herbivores has been well documented, such as reduced elk population density in Yellowstone National Park leading to increased aspen growth [8, 17]. However, the physiological and functional plant traits that herbivores impact can be species-specific rather than general across taxa. Here we have shown a correlation between browsing pressure and leaf architecture via leaf and stem vascular trait responses that appear species-dependent and coordinated with leaf development. We also found that browsing generally led to a decrease in xylem capacity, although by different pathways for each species. Most surprising was the coordination and scaling effects observed in leaves instead of an alteration of anatomical structure, resulting in browsed leaves with stomata and epidermal cell traits comparable to non-browsed leaves. There appears to be a strong species-specific response to herbivory occurring within this system, where browsing impacts functional traits in alternate ways across different species. Future studies should consider the species-specific responses to herbivory to help elucidate the mechanisms of coordinated trait development and especially consider additional factors that were beyond the scope of this study, such as the timing of herbivory in relation to phenology, the inclusion of multiple herbivore interactions on leaf and stem development, and the interplay of environmental factors such as soil moisture climate that could impact response to browsing [48, 49, 50].

## Acknowledgments

A sincere thank you to Alex Baer, Pittermann Lab Manager, for technical assistance and to Alex Jones, UCSC Campus Natural Reserve Manager, for access to campus land.

## Notes

### Competing Interest Statement

The authors have declared no competing interest.

